# Mechanistic insights into the deleterious role of nasu-hakola disease associated TREM2 variants

**DOI:** 10.1101/705608

**Authors:** Raju Dash, Ho Jin Choi, Il Soo Moon

## Abstract

Recently, critical roles of genetic variants in Triggering Receptor Expressed on Myeloid cells 2 (TREM2) for myeloid cells to Alzhimer’s disease have been aggressively highlighted. However, little studies focused to the deleterious role of Nasu-Hakola disease (NHD) associated TREM2 variants. In order to get insights into the contributions of these variants in neurodegeneration, we investigated the influences of three well-known NHD associated TREM2 mutations (Y38C, T66M and V126G) on the loss-of-function by using conventional molecular dynamics simulation. Compared to the wild type, the mutants produced substantial differences in the collective motions in the loop regions, which not only promotes structural remodelling in complementarity-determining region 2 (CDR2) loop but also in CDR1 loop, through changing the inter and intra-loop hydrogen bonding network. In addition, the structural studies from free energy landscape showed that Y38, T66 and V126 are crucial for maintaining structural features of CDR1 and CDR2 loops, while their mutation at this position produced steric clash and thus contributes to the structural impact and loss of ligand binding. These results revealed that the presence of the mutations in TREM2 ectodomain induced flexibility and promotes structural alterations. Dynamical scenarios, which are provided by the present study, may be critical to our understanding of the role of the three TREM2 mutations in neurodegenerative diseases.

## Introduction

Accumulation of amyloid-β (Aβ) in brain parenchyma is the main hallmark of Alzheimer’s disease, which is implicated to the slowly progressive decline of cognitive function, as a result of intracellular neurofibrillary tangles formation, synapse loss and cell death ^1,2^. Although the precise mechanisms and molecular determinants of this neurodegenerative disease remain incomplete, recent studies covering with whole genome sequencing identified that the altered genetic loci including Triggering Receptor Expressed on Myeloid cells 2 (TREM2) is correlated with a drastically increased risk of progression to Alzheimer’s disease ^3,4^.

TREM2 is expressed especially in osteoclasts, microglia, alveolar macrophages, and other mononuclear phagocytes, which is a V-type immunoglobulin (Ig) domain-containing transmembrane protein ^4^. The activation of TREM2 is initiated by anionic lipid such as bacterial lipopolysaccharide (LPS) and phospholipids, and also several putative ligands like apolipoprotein J (apoJ) and apolipoprotein E (apoE) are reported to bind with TREM2 ^5–7^. Upon the binding of ligands, TREM2 recruits protein tyrosine kinase SYK through an adapter protein, which is associated with the transmembrane region of TREM2, is known as DNAX-activating protein of 12 kDa (DAP12) ^8–10^. Subsequently, a cascade of signaling events, including MAPK and PI3K activations is followed and thus regulates the phagocytosis of cellular debris and inflammatory responses of microglia ^10–12^. In both early and mid-term AD, TREM2 plays a protective role ^13^, where its overexpression is involved to clearing the soluble and insoluble Aβ42 aggregates from the brain ^14,15^. On the other hand, TREM2 is reported to avert the accumulation and diffusion of Aβ by modulating microglial activation around amyloid plaques ^16–19^.

At molecular level, the genetic variations in TREM2 are linked with the nasu-hakola disease (NHD) and frontotemporal dementia, where NHD is characterized by demyelination, early-onset dementia, and bone cyst lipoma and linked with the ectodomain Y38C, T66M, and V126G mutations ^20–22^. According to findings of *Kober et al* ^23^ it has been appeared that the NHD mutations may cause the changes of buried residues in TREM2, and promote misfolding with inconsistent impact on surface expression and aggregation. Previous studies explained the influence of TREM2 structure and function by AD associated variants including R47H. However, the deleterious role of NHD-associated mutations is still need to be studied.

The present study considered Y38C, T66M, and V126G mutants of TREM2 in order to characterize the dynamic behavior of TREM2 protein in wild and mutated state by classical molecular dynamics simulation.

## Results and Discussion

### Wild and Mutant type of TREM2

The natural human TREM2 structure is composed of 230 amino acids, a polypeptide chain that consists of three distinct regions including N-terminal mature ectodomain (ECD, residues 19-174), membrane spanning region (residues 175-195) and C-terminal cytosolic tail (residues 196-230). The rest of amino acids, especially residues 1-18 act as the signaling peptide in TREM2 signaling cascade. As shown in **Figure 1**, the tertiary structure of TREM2 ECD domain is mainly composed of nine β-strands (βA - βF), including three major complementarity-determining regions, dubbed as CDR loops, mainly CDR1 (residues from Pro^37^ to Arg^47^), CDR2 (residues Thr^66^ to Arg^76^) and CDR3 (His^144^ to Glu^117^), respectively. Previous studies showed that CDR2 maintain a stable conformation in normal condition by maintaining hydrogen bonding with the CDR1 loop, which is necessary for ligand interactions ^24^. However, the hydrogen bonding network is lost by R47H variants, as it is involved in the network maintenance, which facilitates conformational remodeling of CDR2 loop. Interestingly, the mutations of our interest, including Y38C and T66M are located in both CDR1 and CDR2 regions, respectively. Therefore, how these mutations contribute to pathological behavior of TREM2 is systematically analyzed in the following subsections.

**Figure 1.**
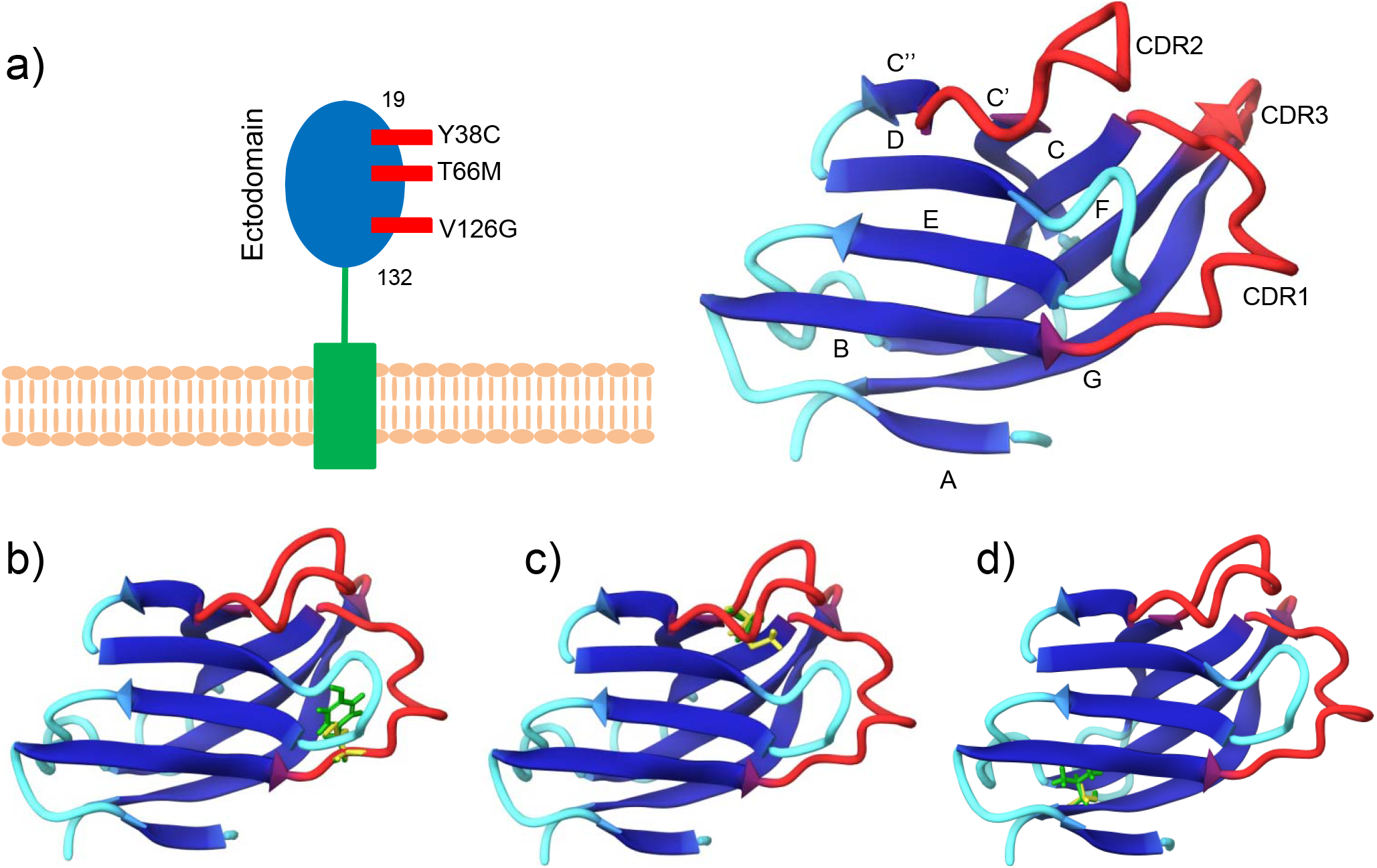
Wild and mutant structures of TREM2 ectodomain. Cartoon depiction of TREM2 wild type ectodomain along with the indication of domain boundaries (a). Three dimensional view of NHD-associated mutated positions, Y38C (b), T66M (c) and V126G (d). Original residue is colored as green, while mutated as yellow.

### Alteration of conformational stability by TREM2 variants

In last decades, the molecular simulation has brought a new platform to detail characterize the structural configuration of macromolecules in various environments at microscopic scale, which is related to their functions and interactions with other molecular species ^25,26^. In this context, molecular dynamics simulations of wild type and other three substitutions (Y38C, T66M and V126G) were conducted for 100 ns of each system to describe their structural dynamics and stability. The conformational stability of the wild and mutant types in the simulation have been analyzed by calculating RMSD for the backbones of all proteins and rendered in **Figure 2**. The RMSD analysis revealed that the wild type and Y38C achieved equilibrium after 5 ns and maintained till to the end of simulation, while rest of the mutants, including T66M and V126G showed system stability following 20 ns. The native and Y38C mutant structures showed similar way of deviation till 100◻ns from their starting structure, resulting in backbone RMSD of ~1.4 to 1.7Å during the simulations. However, T66M and V126G mutant structures showed significantly different deviations from the wild and Y38C structures, ranging from ~1.9 to 3Å. Since the magnitude of fluctuations of four different systems after relaxation period were maintained stable over the time, the simulation is considered to attain convergence, i.e. a stable conformation ^27^, that may lead to produce stable trajectories which is appropriate basis for further analysis. Therefore, the conformational changes in the wild and mutant structure were evaluated by the radius of gyration (Rg) **(Figure 3)**, which describes the overall dimension of protein structure in terms of compactness. Higher values in Rg represent the lower compactness ^28,29^. As shown in **Figure 3 a & b**, the overall dimension of Y38C and T66M structure maintained similar behaviour as like wild type structure, while V126G (**Figure 3c**) structure showed increased compactness after 50 ns, till to the end of simulation. The SASA, which account for solvent accessibility, showed similar profile like Rg for mutants and wild structure. Whereas native and mutant (Y38C and T66M) structures showed similar fashion of deviation from the initial structure (**Figure 4 a & b**), other than V126G structure increased the SASA after 50 ns (**Figure 4c**). Increase in SASA of mutant denotes its relatively bloated nature as compared to the native structure ^30^. Although RMSD analysis depicted the alterations in TREM2 conformation by T66M and V126G mutations, V126G alone showed structure alteration by means of Rg and SASA, which concluded that this mutation decreased the overall dimension of the protein that leads to the increase in solvent accessibility in order to destabilized the protein folding and affecting protein-protein interactions ^31^.

**Figure 2.**
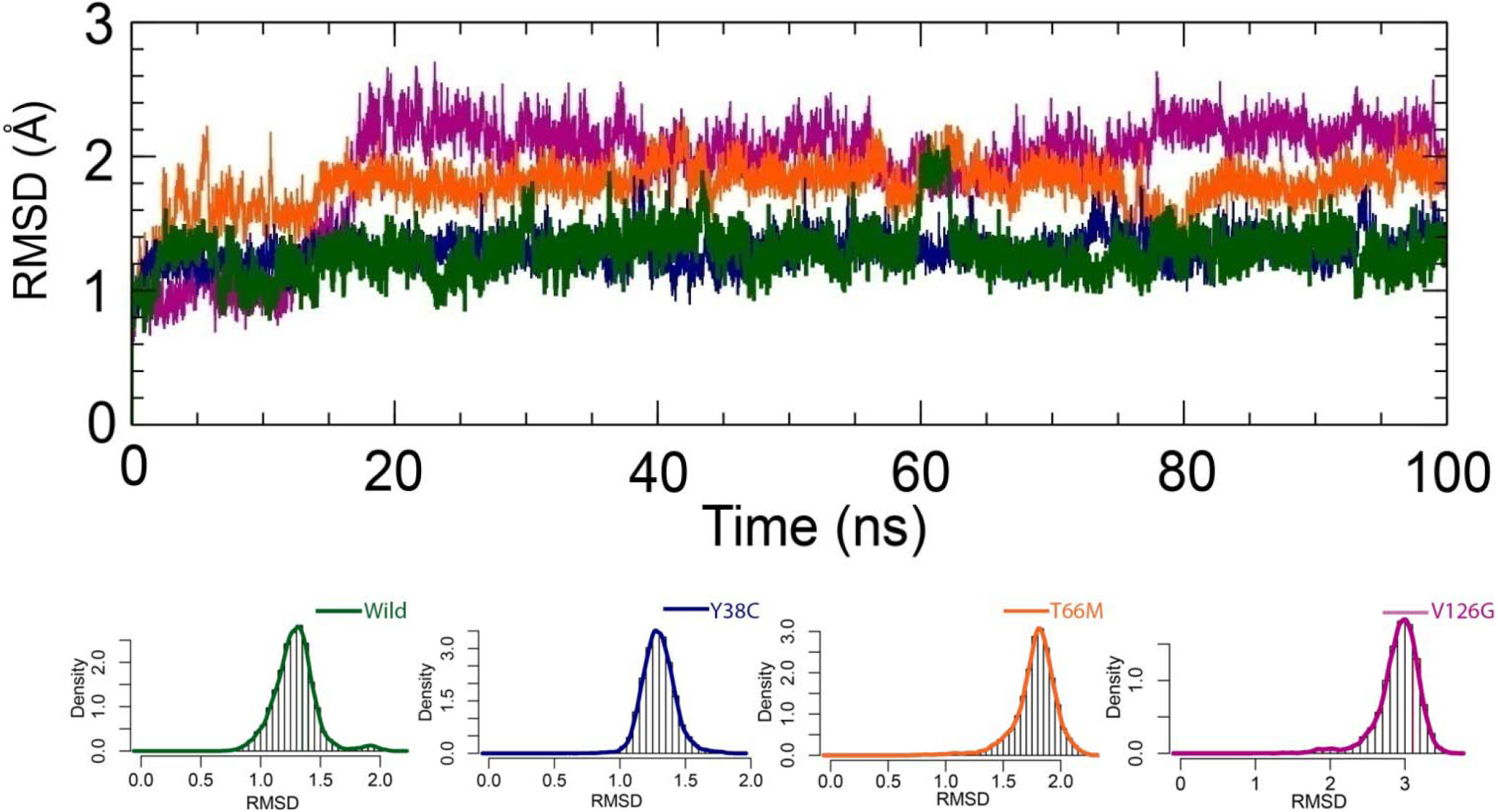
Root-mean-square deviation (RMSD) of the C_*α*_ atoms for the wild-type and mutant TREM2 structure at 100 ns. Here, dark green line represents the wild-type, while the blue, orange and violate lines stand for the Y38C, T66M and V126G mutants of RMSDs. In addition, bottom panel of the figure is represented with the density plots for both wild and mutant structures, illustrating the distribution of sampled conformations during simulation.

**Figure 3.**
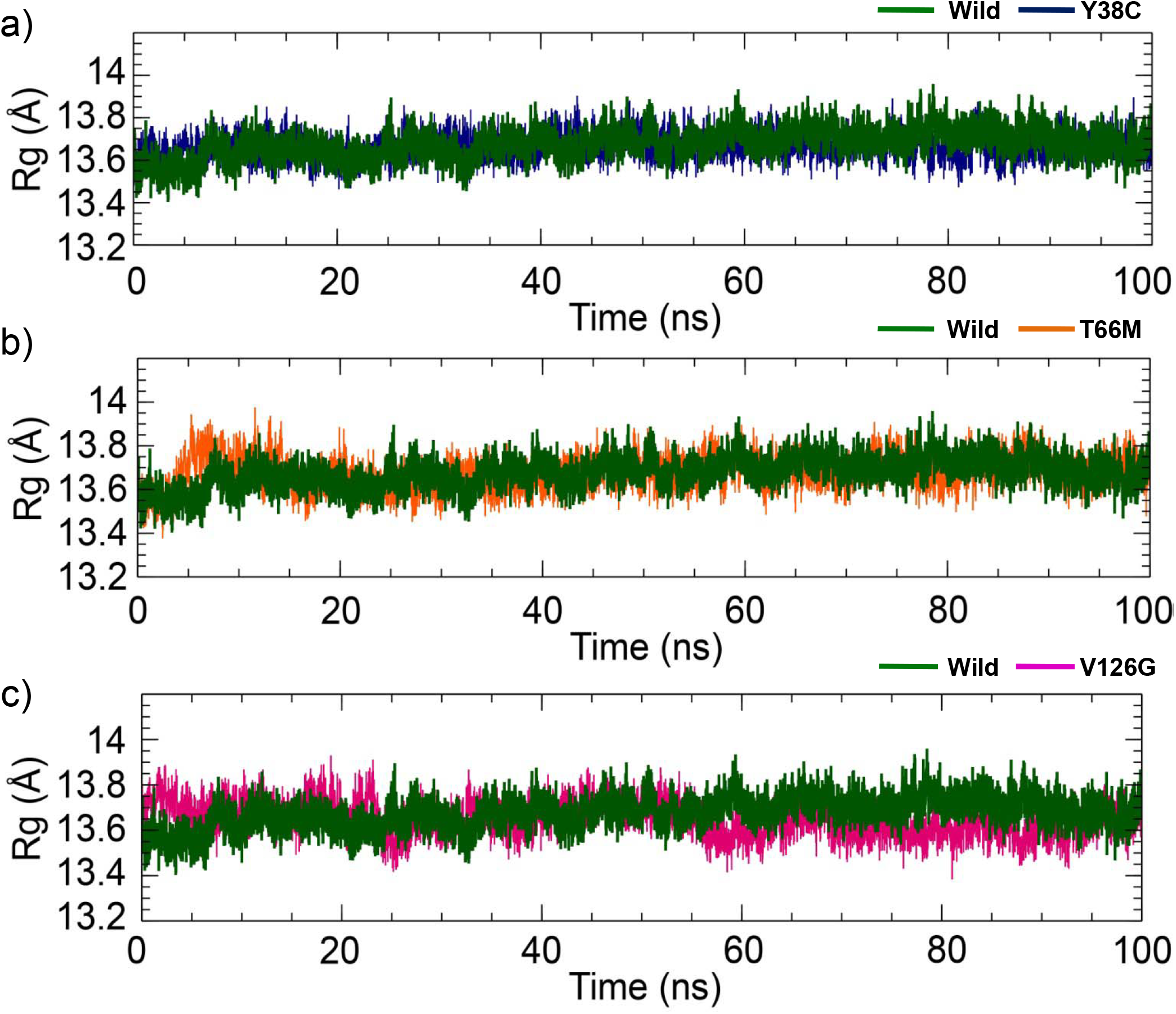
The radius of gyration for the Y38C (a), T66M (b), and V126G (c) TREM2 mutant in comparison to wild type structure. Here, dark green line represents the wild-type, while the blue, orange and violate lines stand for the Y38C, T66M and V126G mutant structure.

**Figure 4.**
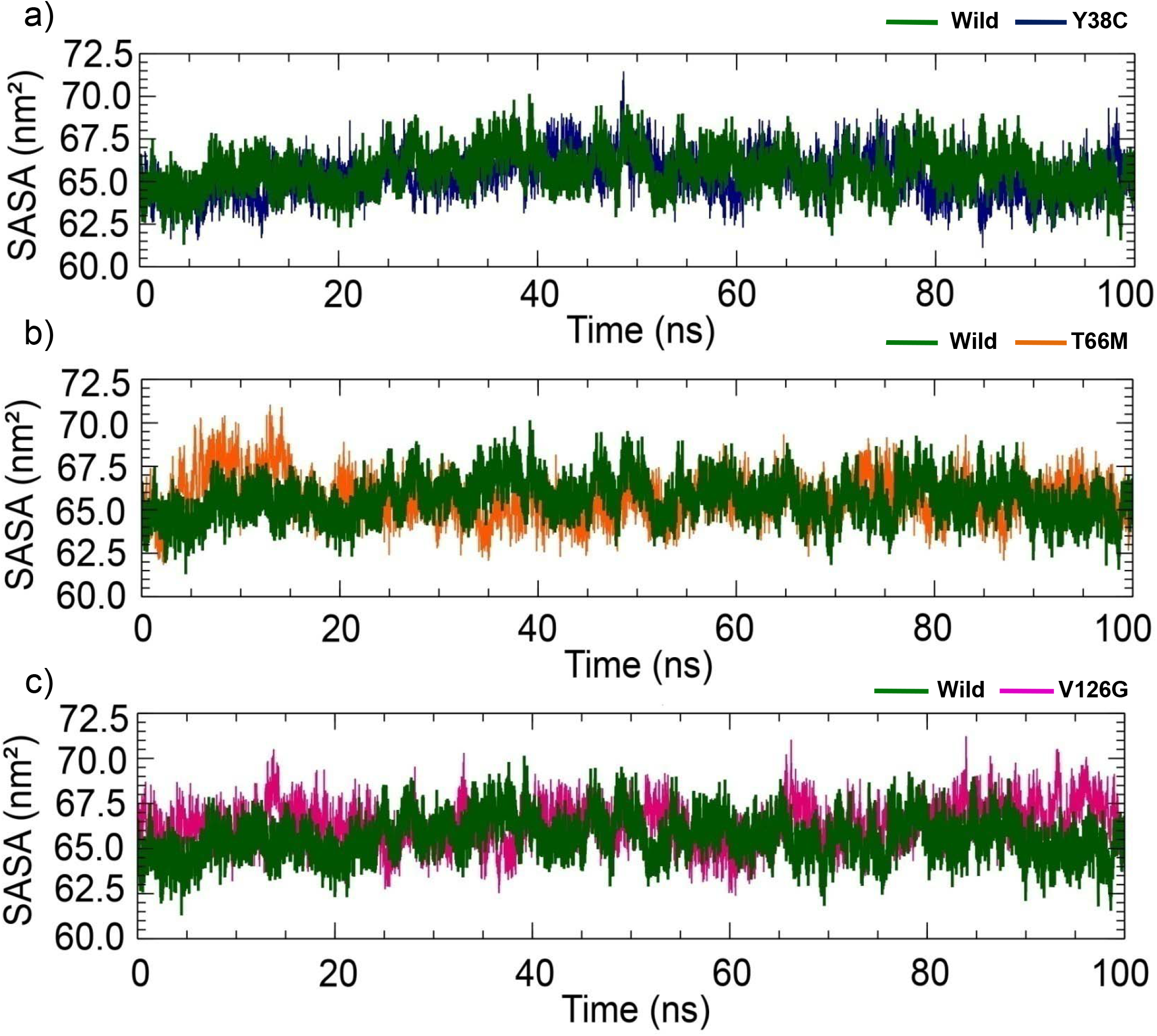
The change of total solvent accessible surface area (SASA) for the Y38C (a), T66M (b), and V126G (c) TREM2 mutant in comparison to wild type structures. Here, dark green line represents the wild-type, while the blue, orange and violate lines stand for the Y38C, T66M and V126G mutant structure.

**Figure 5** rendered RMSF of the each protein backbone, which shows the local changes in protein structure due to the mutations. The local residue fluctuations in the protein serve as an important factor to describe the biological functions, as the many functional sites in the protein have the structural fluctuations and uniquely coupled ^32–34^. As represented in **Figure 5a**, TREM2 structure contains ten loops, including three CDR loops; all were mostly fluctuated throughout the simulations. The degree of fluctuation is high in mutant structures than the wild type. Interestingly, Y38C and T66M mutant structures showed high fluctuations in the CDR1 and CDR2 regions, respectively. On the other hand, V126G mutant increased overall local flexibility of the protein, especially in the regions including residues 50 to 60 and 85 to 95, CDR1 loop and N-terminal tail, respectively. This analysis suggested that mutations have significant impact on the local structural rearrangement rather than overall structure. However, to view more insight, per residue SASA results of all wild and mutant structure have been calculated and represented in **Figure 5b**. The mutant structures increased local solvent accessibility of the protein, especially by T66M in CDR1 and CDR2 regions (**Figure 5b**). Comparatively, the deviations in SASA were also observed by the Y38C and V126G mutants, and it was noticed in the CDR1 and CDR2 loop regions. The SASA of per residue elucidates the structural conformational changes to the protein surface, where lower value means thermodynamically stability as well as buried inside the core of the protein ^35–37^. This analysis together with RMSF study suggested that nasu-hakola disease-associated TREM2 variants provide larger surface area than the native, which is thermodynamically unstable and have major impact on ligand binding.

**Figure 5.**
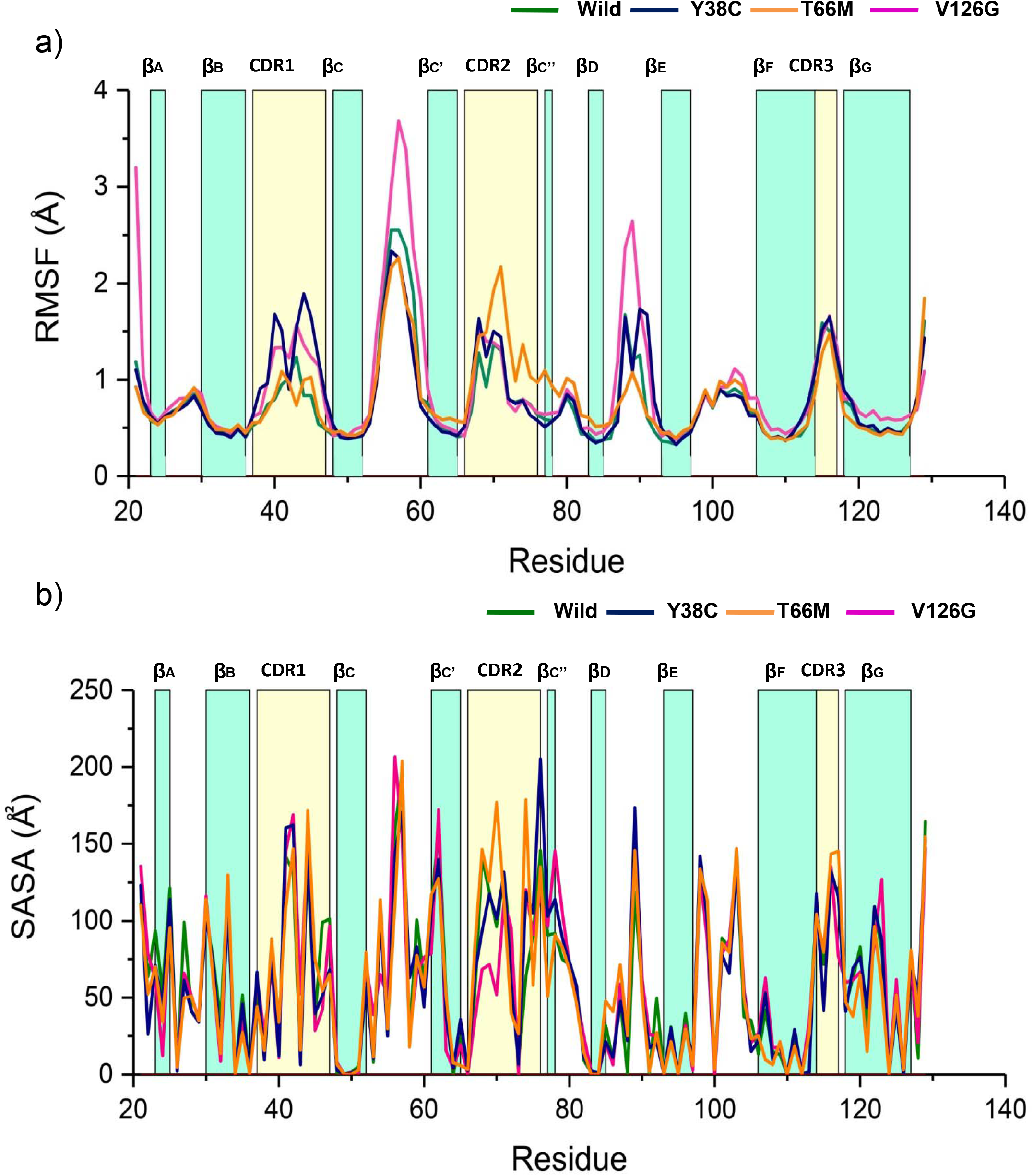
Investigation of local structural effect of mutations. a) Root mean square fluctuation of the C_α_ atoms for the wild type and mutant types of TREM2. a) The SASA values, that computed on a per residue basis for the wild type and mutant types of TREM2. In all plots, residues index is level with colored bar, representing the structural propensity. In each graph, dark green line represents the wild-type, while the blue, orange and violate lines stand for the Y38C, T66M and V126G mutant structure.

### Effect of mutations in protein dynamics

In different functional states, protein undergoes various conformational transitions by the global domain motions, which is facilitated by the collective motion of backbone atoms ^38^. Principally, various regions including the regions having secondary structures move in a correlated manner ^39^. For additional insights into concerted dynamics of TREM2 in wild and mutant types, DCCM and principle component analyses were performed on the MD trajectories from different system. The DCCM covers total correlated motions among the protein residues, where the strong correlated motions between specific residues are highlighted by red regions (Figure 6). On the other hand, highly anti-correlation is expressed by blue regions. The analysis revealed that mutations induced completely different motions than the wild type structure. Comparatively, Y38C (**Figure 6b**) produced significant anti-correlative motion between residues 17-27 (CDR1 loop, residues 37 to 47) and residues 45 to 56 (CDR2 loop, residues 65 to 76) as well as 65 to 74 (residues, 85 to 94). On the other hand, T66M (**Figure 6c**) slightly reduced the degree of both correlated and anti-correlated motions that exist in the wild type structure, although no significant correlation has been seen in the DCCM analysis. In case of V126G mutant structure (**Figure 6d**), the result showed that the mutation increased the correlative intra-residue movement, specifically between the loop 34 to 40 (residues, 54 to 60) and c-terminal portion 80 to 105 (residues, 100 to 125). The negative correlation was also seen in the CDR1 and CDR2 loop regions, indicating that the deleterious effect in TREM2 functions.

**Figure 6.**
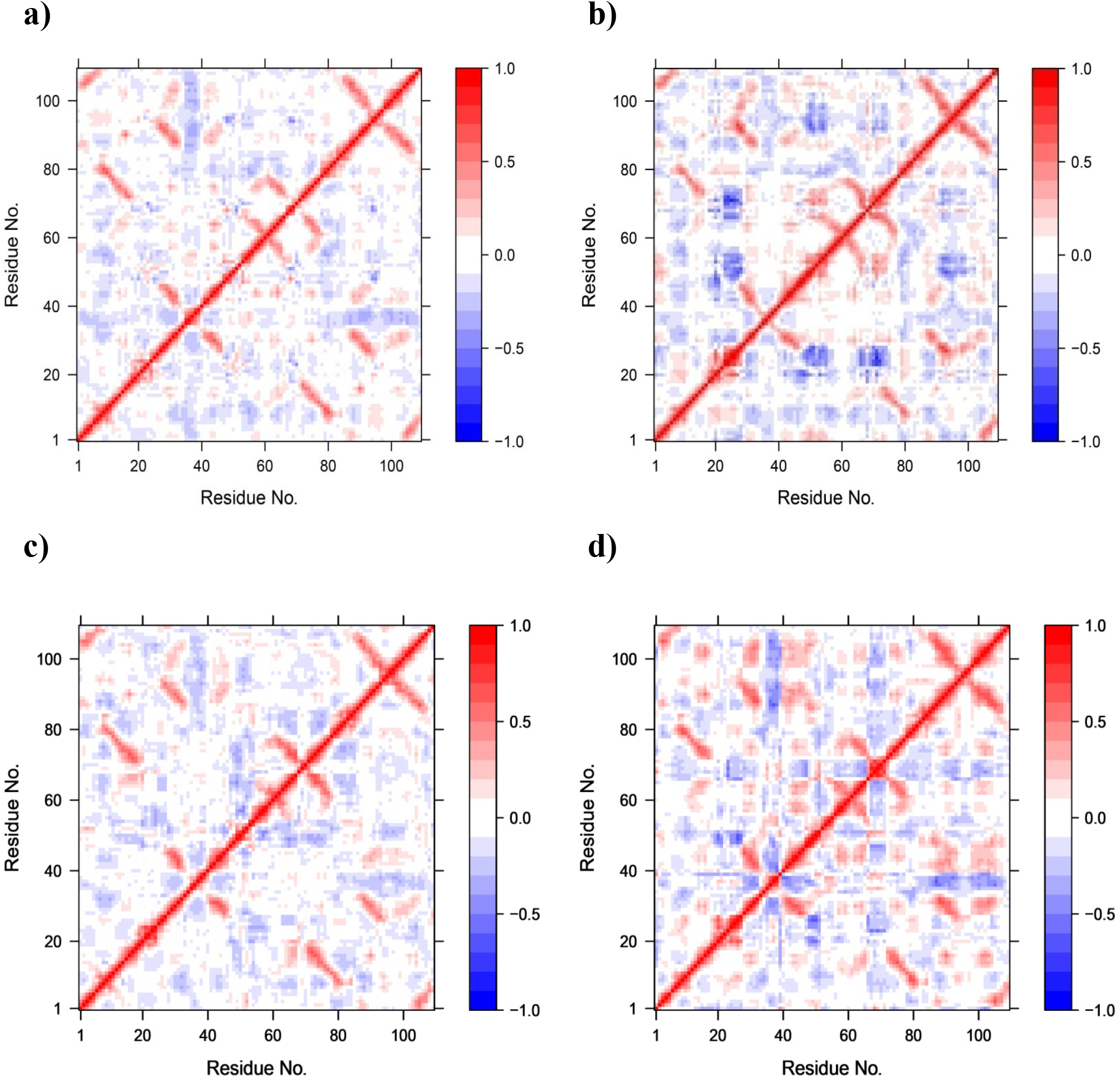
Calculated dynamic cross-correlation matrix of the C_α_ atoms around their mean positions for 100◻ns molecular dynamics simulations. The extent of correlated motion and anti-correlated motions are color-coded, ranging from red to blue. Here red color denoted positive correlation, while negative described by blue color. Wild type (a), Y38C (b), T66M (c) and V126G (d).

To further support this explanation, principal component analysis (PCA) was conducted based on C_α_ atoms (**Figure S1**). In order to get insights about the dynamic mechanical properties of the simulated system, PCA analysis is usually performed ^26^, by reducing the complexity of classifying collective motions. The dynamics of two different proteins is best achieved via characterization of its phase space behavior, which is directly associated with the protein stability and protein function ^40,41^. At this point, the overall combined motion of the C_α_ atoms of protein structures were described by the eigenvectors of the covariance matrix, which is correlated with the coincident eigenvalues. As resulted in the PCA analysis, the simulated systems including, wild type, Y38C, T66M and V126G showed 19.81%, 33.43%, 20.94% and 29.67% of variance in the first three PCAs, respectively.

The motions of different proteins can be visualized through the projection of trajectories in phase space of the first two principal components (PC1, PC2), where mutants show different patterns of conformational spaces, denoting the significant changes in protein conformation. On these projections, the Y38C showed uniform conformational distribution in a larger phase space, describing the higher flexibility in contrast to the native (**Figure S1b**). Furthermore, T66M structure also showed conformational transitions in the simulation, representing the less flexibility over the time (**Figure S1c**). On the contrary, V126G represented higher periodic jumps, indicating greater fluctuations compared to the native, as it occupied a larger phase space (**Figure S1d**).

The variations in atomic movement were represented by all first PCAs (**Figure 7)**, as PC1 reveal the most dynamic motion of all systems. Here the displacement of atoms is denoted by wide tubes, whereas the narrow tubes marked the regions that stayed rigid during the simulation. The structural deviations among the wild type and mutations were clearly depicted in the first PCAs (**Figure 7)**, where the contribution of each residue to the first principal component is represented in the plot which is rendered in the bottom panel (**Figure 7e**) describing the local mobility during the simulation. As can be seen, Y38C induced flexibility in several loops of TREM2 including CDR1, CDR2 and CDR3 regions (**Figure 7b**), while CDR2 was greatly fluctuated in T66M mutant structure (**Figure 7c)**. The conformational changes in the wild type structure were reflected in the loop region, residues 47 to 60; this region also deviated in V126G mutant structure (**Figure 7d)**. The PCA analysis supports the results described by the RMSF and DCCM analysis, which ultimately concluded that the mutants have higher structural flexibility than the wild type demonstrating local deleterious effects. Since Y38C and T66M TREM2 produced large flexibility in CDR regions, these mutations may bring different structural orientations in these regions. The following section further addresses on this issue.

**Figure 7.**
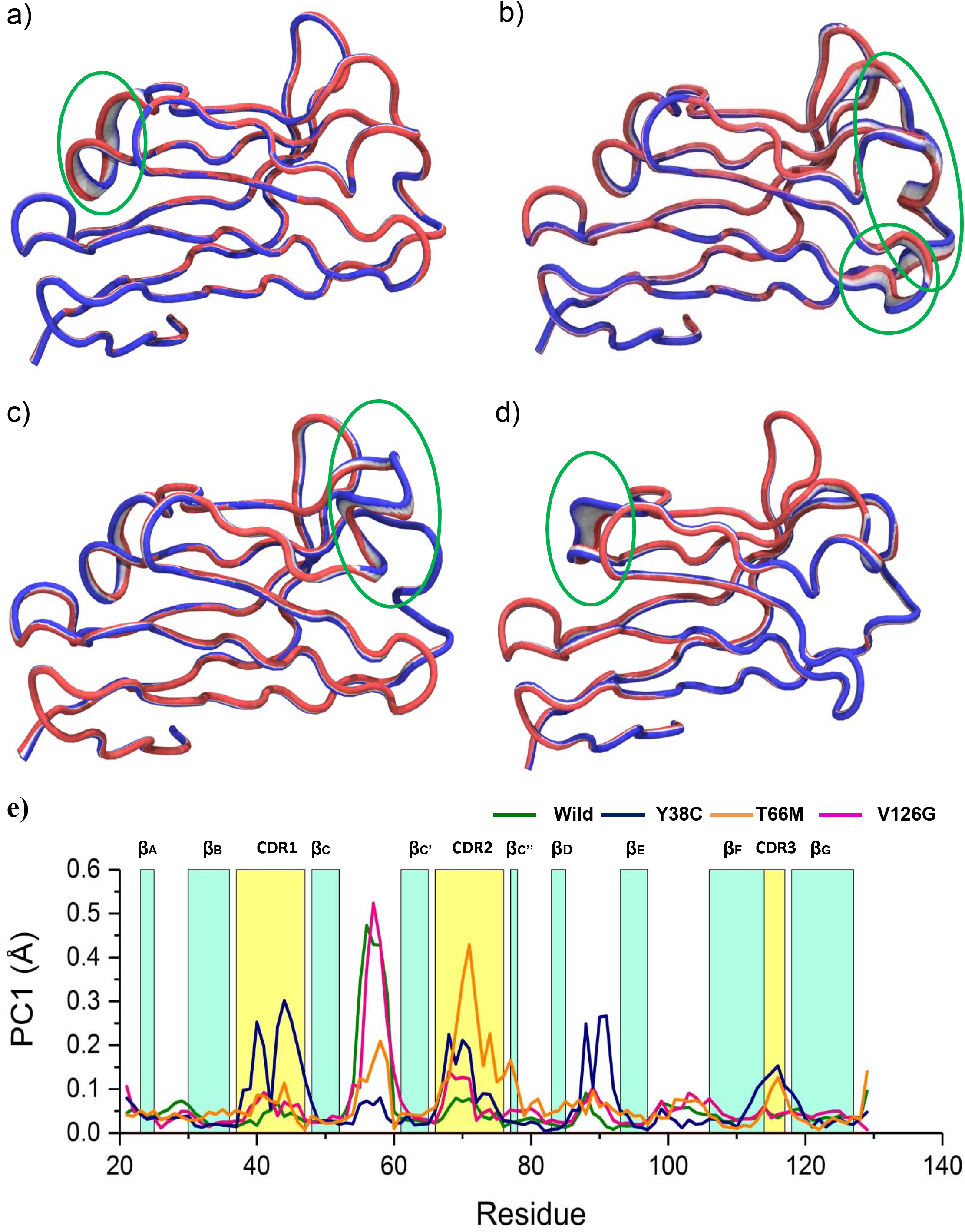
Representation of the atomic displacements in PCA1, calculated from the simulation of wild type (a), Y38C (b), T66M (c) and V126G (d), which indicates significant alterations in the movements. The green marked circles of each panels describes the regions of greater flexibilities. Residue-wise loadings for PC1, where dark green line represents the wild-type, while the blue, orange and violate lines stand for the Y38C, T66M and V126G mutant structure (e).

### Mutation-induced conformational changes

Although MD simulations well explained the conformational changes in the protein structure, the changes in the secondary structures caused by the mutations are still needed to investigate. Therefore, the variations in the secondary structural elements for both wild and mutant proteins during the simulation were characterized by the simulation interaction diagram of Schrödinger 2017-1 (LLC, New York, NY, USA) ^42^. During the simulation, the appearance of secondary structural elements determines the structural flexibility of a protein molecule, where conformations such as α-helix and β-sheet are naturally tend to be more rigid than the coils and turn conformations ^26^. The total percentage of secondary structural elements in total simulation time is shown in **Figure S2**. Interestingly, the mutants increased the total secondary structures of TREM2 protein, when compared to wild type. Moreover, the CDR1 (residue index 17-27) and CDR2 (residue index 46-57) regions were grained short α-helix conformation in the mutant structure, where the percentage of helix formation is higher in the Y38C structure (**Table S1**). On the contrary, V126G increased total secondary structure elements in the TREM2, where the increase of α-helix conformation in CDR2 region was higher in comparison to other mutants. Previous studies with Alzheimer’s disease associated mutant showed that substation of ARG^47^ by His facilitates CDR2 loop remodelling to short α-helix which contributes to loss of ligand binding ^24^. Together with these data, our study also suggested that mutants in this study also facilitate not only conformation remodelling of CDR2 loop but also in CDR1 regions.

### Mechanistic insights into conformational remodelling

It is inferred that the intra-residue hydrogen bonding influences the secondary structure in biological macromolecule ^43^. Thus hydrogen bond occupancy analysis within CDR1 and CDR2 loops in the wild as well as in mutant types was further considered, as both CDR1 and CDR2 loops undergoes conformational remodelling (**Figure S2**). As shown in the **Figure 8**, mutations induced diverse intra and inter loop hydrogen bond interactions. Furthermore, these loops were identified as the flexible regions from RMSF and PCA studies, indicating that increasing flexibility by mutations render residues to make different interaction network. According to the RMSF study, the substitution of tyrosine with cystine in 38^th^ position was seen to induce high flexibility next to D39, which formed significantly elevated hydrogen bonding interactions within CDR1 loop, especially with M41 and H43 (**Figure 8a**). The hydrogen bonding occupancy between D39 and H43 in Y38C structure was maintained to 12.34% during simulation, while it was 3.91% in wild type. Furthermore, D39 maintained 6.6% H-bond occupancy with M41 residue; however, it was missing in wild type structure. This observation indicates that maintaining intra-loop hydrogen bond with M41 and H43 by D39 may facilitate short helix formation of CDR1 loop in Y38C mutant. In case of CDR2 loop, Y38C mutant reduced hydrogen bonding between S73 and W70, L72 and S75, while it was increased in the wild type. Besides, Y38C maintained two new hydrogen bonds, including T66 and S73, S73 and L69, which were lacking in wild type structure (**Figure 8b**).

**Figure 8.**
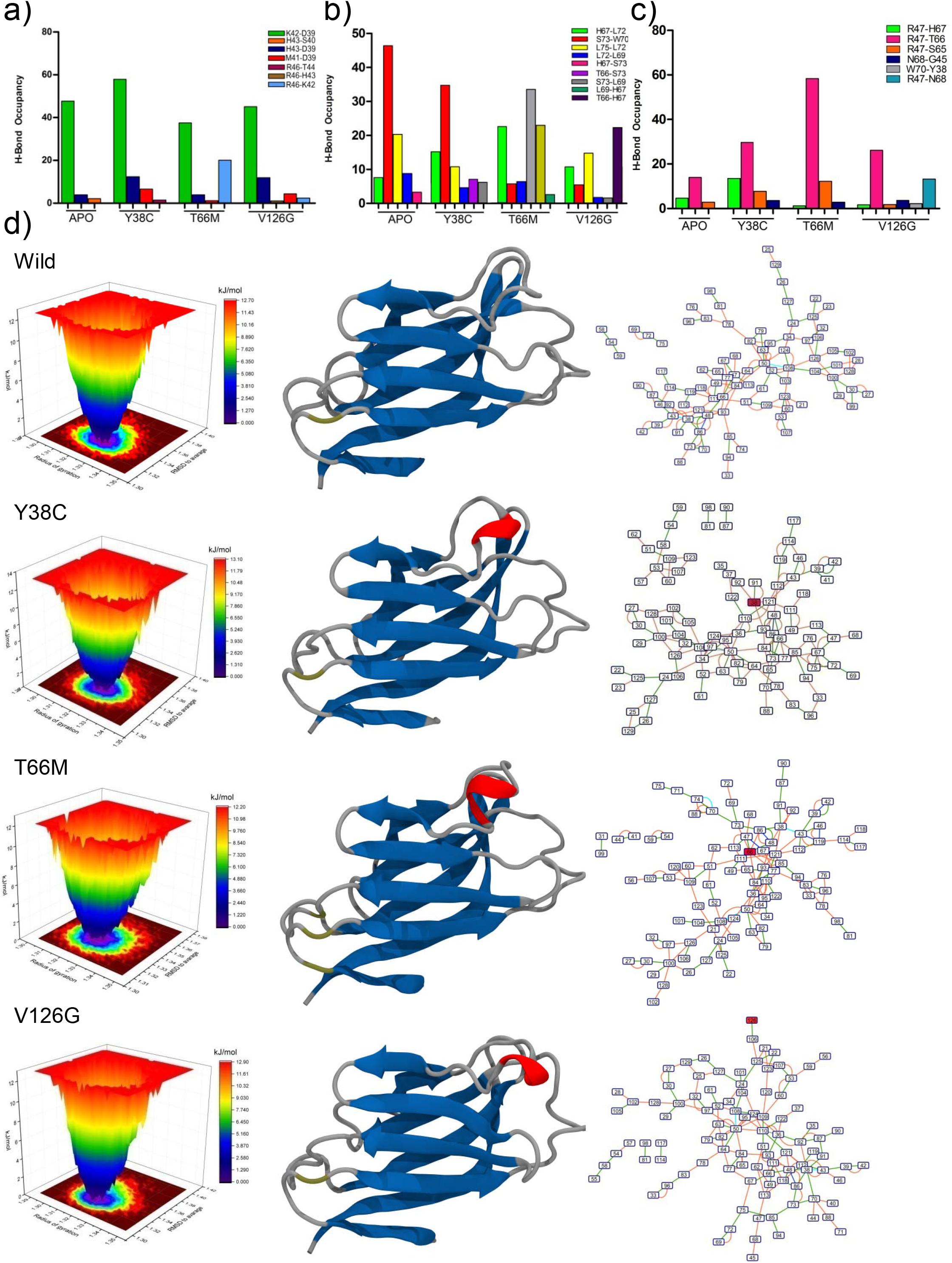
Hydrogen bond occupancy analysis for within the CDR1 (a), CDR2 (b), between the CDR1 and CDR2 loop (c). Free energy contour map depicted based on the Radius of gyration and RMSD to average (d), where deeper dark blue color area in the maps indicates lower energy. In the figure each graph is represented with the three dimensional structure having the lowest energy, along with residue interaction network. Each node in network describes the residue, while edge represented as the interaction. The color of edge signifying the mode of interaction, including green for hydrogen bond, orange of van der walls contact, cyan color for salt bridge, pi-pi stacking as blue and pink for disulfide bridge.

Previous studies showed that the conformational change of CDR2 loop is facilitated by the π-π stacking interactions between the imidazole moieties of H47 and H67, which caused to swing H67 approximately by 180° ^24^. Therefore, inter-loop interaction between CDR1 and CDR2 was investigated and found that Y38C increased the hydrogen bonding between H47 and H67 (**Figure 8c**). This observation indicates the conformational changes in CDR1 loop. Similarly, T66M mutant showed the significant local fluctuations in S65 and H67 residues, compared to wild type; H67 maintained H-bonding interaction with L72 more than 20%. Interestingly, mutation in T66 increased the hydrogen bonding capacity with R47 more than the wild structure. However, an unusual hydrogen bond interaction between R47 and D65 by replacing M66 was seen, which may contribute to CDR1 conformational change. However, the interaction between M66 and R47 was more dominant, as a result the degree of helix formation in CDR1 loop was seen to decrease subsequently in the simulation (**Figure S2c**). On the other hand, V126G mutation reduced the hydrogen bonding between R47 and T66, while induced interaction between R47 and N68, which might allows the conformational changes in CDR1 loop. In addition, conformational changes in CDR2 loop could be induced by numerous hydrogen interactions of H67 with M41, H43 and T66.

In order to validate the findings from H-bond analysis, additional free energy landscape analysis was introduced to select most stable conformer from the simulation ensemble. The FEL is rendered in **Figure 8d**, where different colors represent the corresponding structures with different energies. The conformation having the lower energy states can be found in the blue area, are usually more stable than the other conformations (red area) generated during the simulation ^44^.

The thermodynamic stability of a protein is represented by the depth of an energy minimum, where the kinetic stability of a protein is indicated by the height of barriers separated by different energy minima. This depth of energy also suggests the change of conformation from one to another, where the width of an energy minimum associated with the width of the conformational ensemble within the energy well ^45^. As represented in the **Figure 8d**, the region of the conformational space corresponding to the basin was seen to change in both wild and mutant types, indicating that the overall conformational stability of the protein was affected by the mutations. In order to understand the disordered state, the conformation in the favorable energy minima has been visualized and analyzed the intra-residue hydrogen bonding network through residue interaction network analysis. The lowest minima in wild type trajectory was found with corresponding Rg and RMSD to average value of 1.33 and 1.32 nm, respectively, containing no conformational changes in CDR1 and CDR2 loops. In detailed structural view, T66 was found to interact with R47 by forming double hydrogen bonds. Comparatively, a short helix was found in the CDR2 loop in the most stable conformer of mutant trajectories. In Y38C mutant structure, M41 was seen to be maintained H-bond with D39. Furthermore, R47 formed hydrogen bonding interactions with both H67 and L72 in Y38C, while in case of T66M; this residue maintained Pi-Pi stacking interaction with H67. On the other hand, V126G structure showed an unusual hydrogen bonding between 66 and 67 residues as well as between R47 and S65. Although no conformational change in the CDR1 loop was visualized in the structure, the interaction between D39 and M43 was also seen within CDR1 loop. Most importantly, all mutants, especially Y38C and T66M, produced steric clashes with neighbouring side chains, while mutation with glycine at 126 position induced side chain steric clash within Y38 residue, which induced flexibility in both CDR1 and CDR2 loops thus promoting conformational remodelling.

## Conclusion

The effect of genetic variants in the loss of TREM2 function is crucial to understand its involvement with late-onset Alzheimer’s disease (AD). Using molecular dynamics simulation, this study represented novel findings on the deleterious role of NHD variants, which promotes structural alteration in TREM2 ectodomain. All three mutations (Y38C, T66M and V126G) induced structural remodelling in both CDR1 and CDR2 loop by inducing steric clash, increased flexibility and altered hydrogen bonding patterns. This study supports previous findings, and provides additional insight on the mechanism behind the loss of ligand binding, which is critical to our understanding of the role of TREM2 in neurodegenerative diseases.

## Methods

### Preparation of simulation system

The three dimensional crystal structure of TREM2 ectodomain was retrieved from the protein databank (http://www.rcsb.org/pdb) ^46^, with PDB id of 5ELI ^47^. The structure was initially prepared by adding bond orders, hydrogen and charges; and also refined by removing water molecules and optimizing it at neutral pH. The structure was further fixed by correcting amide groups of asparagines, some thiol and hydroxyl groups, protonation states of glutamic acids, aspartic acids and histidines. In order to adjust the heavy atom Root Mean Square Deviation (RMSD), minimization was applied by using Optimized Potentials for Liquid Simulation (OLPS3) force field until it reached to 0.30 Å. The mutant structures, consisting of Y38C, T66M, and V126G, were constructed by computational mutagenesis methodology, using the Mutate Residues script from Schrödinger suite 2017-1 (LLC, New York, NY, USA) ^48^.

Additional short molecular dynamics refinement simulation was performed for better resolute quality of the structure. Thus applying YAMBER3 force field ^49^, molecular dynamics simulation was conducted for 500 ps at pH 7.4 and 298K temperature within the solvent density of 0.997. The simulation was run by the YASARA software with a default md_refine macro, and the final structure was selected based on the lowest energy.

### Molecular dynamics simulation

Molecular dynamics simulation approach was used to study the changes in dynamics behavior of protein due to mutations, by the implementing Desmond module of Schrödinger suite 2017-1 (LLC, New York, NY, USA) ^50,51^. Here, OLPS3 all-atom force field was utilized to illustrate the molecular behavior ^52–54^. The prepared structures including wild type, Y38C, T66M, and V126G were solvated in the presence of explicit solvent in a triclinic periodic boundary box. Each systems was submerged to a Monte-Carlo equilibrated TIP3P solvation model extending to approximately 10Å in each direction, which was known to give the best experimental output ^55^. For the neutralization of the solvation system, additional counter ions were added to the water model. The physiological condition as well as the ionic strength of the solvent system was also maintained by placing the default salt (NaCl) concentration of 0.15◻M to the simulation box. The default relaxation protocol ^56,57^ was used to relax the system, which contains eight stages. Followed by Brownian dynamics, the second stage simulation was started at the temperature of 10K in NVT ensemble for 12 ps and restraints on solute heavy atoms. Afterward, 12 ps simulation was done in same ensemble at temperature of 10K and restraints on solute heavy atoms in third stage. In the same temperature, the fourth stage was begun by running 12 ps simulation under the NPT ensemble at 1 bar pressure by keeping restraints on the solute heavy atoms. The protein cavity in the stage five solvated by using the solvate pocket script. Followed by 12 ps simulation in NPT ensemble in stage six with restraints on the solute heavy atoms, 24ps simulation was run in the stage seven with no restraints on the solute heavy atoms. Both stages were performed at the 300 K in 1 bar pressure. Finally 100 ns MD simulation was done for each system with no constraints applied and resulted trajectories were subjected for further analysis. Throughout the simulation, the integration of motion, RESPA ^58^ integrator was employed with 2 fs as inner time step with and M-SHAKE ^59^ algorithm to constrain all covalent bonds connecting hydrogen atoms. Particle mesh Ewald method was applied for calculating the long-range electrostatic interaction [22], while 9.0 Å was used for short-range electrostatic contacts; uniform density approximation was select for the cutoff of the long-range van der Waals (VDW) interaction cutoff. The conditions in the simulation were maintained by Nose–Hoover thermostats ^60^ at 300 K constant temperature and 1 atmosphere constant pressure using the Martyna–Tobias–Klein method ^61^.

The resultant trajectories were used to evaluate the conformational change and protein stability by means of Root Mean Square Deviation (RMSD), Root Mean Square Fluctuation (RMSF) and SSE (Secondary Structure Elements) by using Simulation Interactions Diagram panel of Schrödinger 2017-1 (LLC, New York, NY, USA). Whereas, other analysis including, radius of gyration (Rg). Solvent Accessible Surface Area (SASA) and hydrogen bond occupancy were calculated by VMD software ^62^. The dynamic cross-correlation maps were generated from trajectories to explain the time-correlated motions in the protein, which were assembled by Bio3D [41] software through R programming. The time-correlated information between protein atoms i and j (c_ij_) is represented as a matrix in DCCM, which was obtained with the following expression:

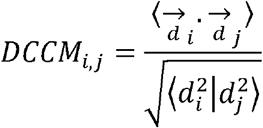

The displacement between the current and average position of _*i*_*th* and _j_th atoms is represented by *d*, and the mean time over the all trajectories is represented by the angle brackets. The calculated values in DCCM ranged between the −1 to +1, denoting negative and positive correlation, respectively. Principle Component Analysis (PCA) was further included in the study to describe the collective motions of TREM2 variants ^39^. The eigenvectors in PCA were calculated by the superimposing atomic coordinates to the reference structure without translational and rotational movements. Each eigenvector displays the mean squared displacements (MSD) of atoms along the corresponding eigenvector, which was associated with an eigenvalue. The mathematical details have been described previously ^63,64^.

### Free energy landscape (FEL)

Free energy landscape is used to map the all possible conformational changes in macromolecules through highlighting their corresponding energy levels, which required to probe the spatial position of interacting molecules in a system ^65,66^. In free energy landscape analysis, Gibb’s free energy is calculated as a function of protein enthalpy and entropy, which describes the protein stability. It also highlights the various conformational states related to protein’s structure-function correlation. In this study, the energy bases of conformational diversity in various TREM2 structures were investigated by the free energy landscapes, using following equation:

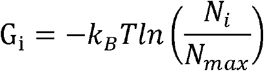

Where, Boltzmann’s constant is represented by *k*_*B*_, *T* defined the temperature, which is set to 300◻K. *N*_*i*_ is the population of bin *i* and *N*_max_ is the population of the most populated bin. An artificial barrier scale is set to the bin with no population, as the lowest provability. Color-code modes are used to display different energy levels.

### Residue Interaction Network

The most stable three dimensional coordinates of wild and mutant types were transferred to RING server ^67^, in order to analyze the Residue Interaction Network (RIN), which represents intraresidue interactions in a exhaustive network view. In the network model, a protein residue is represented as the node, while the mode of interactions is described as the edge. The result from RING server is then subjected to Cytoscape for constructing the interactive RIN 3.2.1 ^68^, using the plug-in RINalyzer. The types of interactions are described by the dashed or dotted edges, including the salt bridge, hydrogen bond van der Waal interactions.

## Author Contributions

ISM conceived the idea and designed the experimental work. RD and HJC carried out the experiments. ISM, RD and HJC analyze the data and wrote the manuscript.

## Supplementary data

**Table S1.** The summery of secondary structure element throughout the protein structure in different simulation systems.

**Figure S1.** Principle Component Analysis (PCA) regarding the TREM2 protein in four different systems, wild type (a), Y38C (b), T66M (c) and V126G (d). Each panel represents the two-dimensional plots between eigenvectors (EV) 1, 2, and 3 in three cross profiles, PC1-PC2, PC1-PC3 and PC2-PC3 for all systems. Throughout the x and y axes, each dot denotes the one conformation of the complex. The spread of blue and red color dots described the degree of conformational changes in the simulation, where color scale from blue to white to red is equivalent to simulation time. The blue indicates initial timestep, white is intermediate and final timestep is represented by red color.

**Figure S2.** Residue-specific average secondary structural preferences for wild type (a) Y38C (b), T66M (c) and V126G (d) structure, during 100◻ns simulation. Here, dark blue color stand for Beta sheets and red color for alpha helix conformation.

