## Supplementary data for "Mechanistic insights into the deleterious role of nasu-hakola disease associated TREM2 variants"

**Table S1.** The summery of secondary structural elements throughout the protein structure in different simulation systems.

| **Name of System** | **% of Helix** | **% of Strand** | **% of Total SSE** |
| --- | --- | --- | --- |
| Wild | 0.40 | 46.97 | 47.37 |
| Y38C | 1.47 | 46.97 | 48.43 |
| T66M | 0.93 | 48.80 | 48.80 |
| V126G | 1.35 | 47.90 | 49.25 |

**Figure S1.** Principle Component Analysis (PCA) regarding the TREM2 protein in four different systems, a) wild type, b) Y38C, c) T66M, d) V126G.

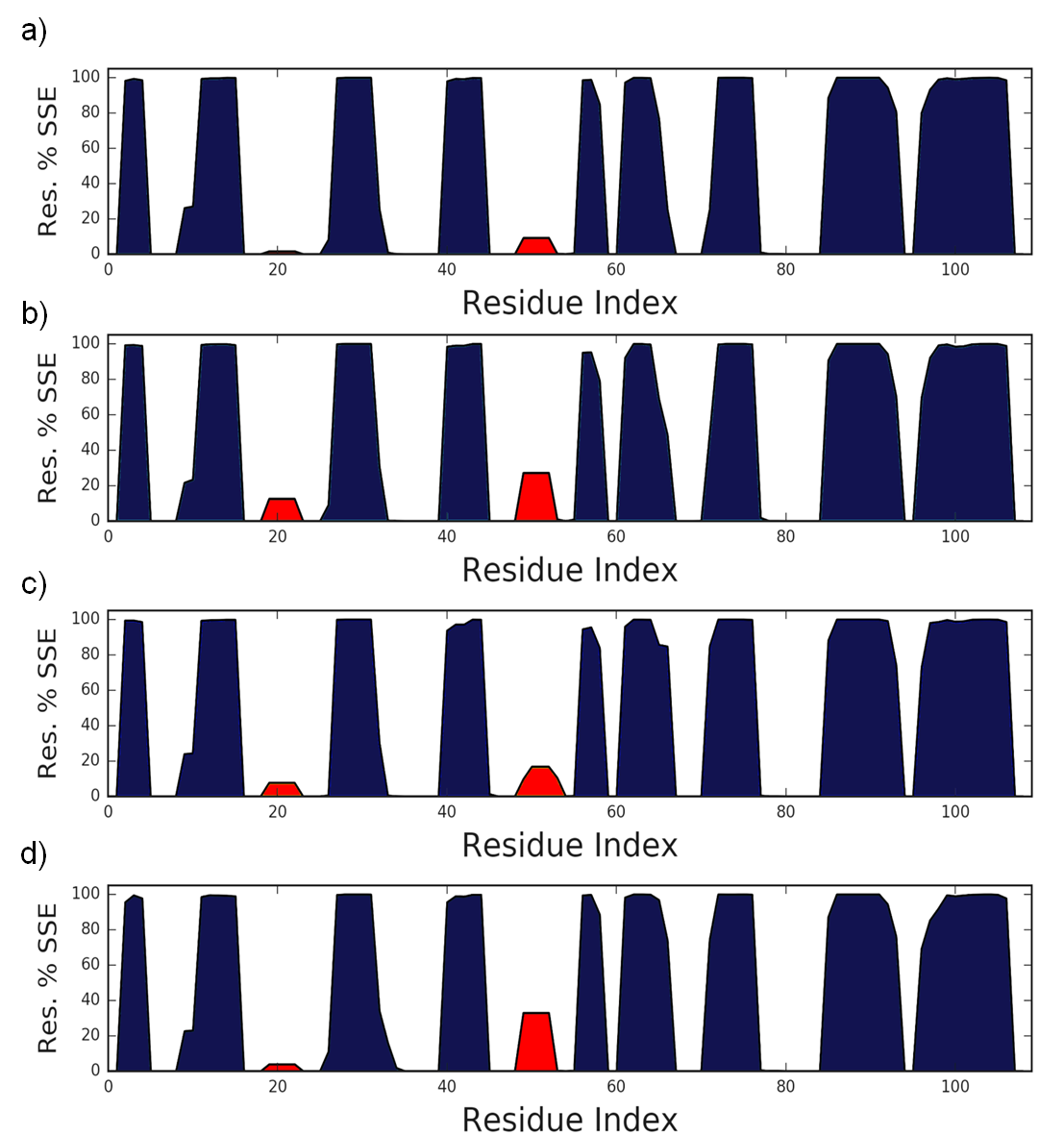

**Figure S2.** Residue-specific average secondary structural preferences for a) wild, b)Y38C, c) T66M, and V126G structure, during 100 ns simulation. Here, dark blue color stand for Beta sheets and red color for alpha helix conformation.
